# Membrane binding of endocytic myosin-1s is inhibited by a class of ankyrin repeat proteins

**DOI:** 10.1101/2023.04.26.538419

**Authors:** Alaina H. Willet, Jun-Song Chen, Liping Ren, Kathleen L. Gould

**Affiliations:** Department of Cell and Developmental Biology, Vanderbilt University School of Medicine, Nashville, TN 37232

**Keywords:** endocytosis, Myo1, plasma membrane, fission yeast, OSTF1, Myo1e

## Abstract

Myosin-1s are monomeric actin-based motors that function at membranes. Myo1 is the single myosin-1 isoform in *Schizosaccharomyces pombe* that works redundantly with Wsp1-Vrp1 to activate the Arp2/3 complex for endocytosis. Here, we identified Ank1 as an uncharacterized cytoplasmic Myo1 binding partner. We found that in *ank1Δ* cells, Myo1 dramatically redistributed from endocytic patches to decorate the entire plasma membrane and endocytosis was defective. Biochemical analysis and structural predictions suggested that the Ank1 ankyrin repeats bind the Myo1 lever arm and the Ank1 acidic tail binds the Myo1 TH1 domain to prevent TH1-dependent Myo1 membrane binding. Indeed, Ank1 over-expression precluded Myo1 membrane localization and recombinant Ank1 blocked purified Myo1 liposome binding in vitro. Based on biochemical and cell biology analyses, we propose budding yeast Ank1 and human OSTF1 are functional Ank1 orthologs and that cytoplasmic sequestration by small ankyrin repeat proteins is a conserved mechanism regulating myosin-1s in endocytosis.

**Summary:** Fission yeast long-tailed myosin-1 binds Ank1. Ank1 ankyrin repeats associate with the Myo1 lever arm and Ank1 acidic tail binds the Myo1 TH1 domain to inhibit Myo1 membrane binding. Ank1 orthologs exists in budding yeast (Ank1) and humans (OSTF1).

## Introduction

Type I myosins are actin binding proteins that link membrane to F-actin to support cellular shape changes. Myosin-Is are characterized by an N-terminal motor domain, a neck region, and a tail region of varying length and composition. The long-tailed subclass of myosin-1s is found in many eukaryotes (Lee *et al*., 2000; Sirotkin *et al*., 2005) and contain extended C-terminal tails that mediate protein-protein interactions, membrane binding, and/or actin binding.

The single type I myosin present in *Schizosaccharomyces pombe*, Myo1, belongs to the long-tailed subclass and localizes to actin patches which assemble at sites of endocytosis (Lee *et al*., 2000). Myo1 contains an N-terminal motor domain followed by a lever arm containing two IQ motifs that bind the calmodulin-related light chains Cam1 and Cam2 (Toya *et al*., 2001; Sammons *et al*., 2011). The C-terminus includes a tail homology 1 (TH1), a tail homology 2 (TH2), a *src* homology 3 (SH3) and a central-acidic (CA) domain. The CA domain binds and activates Arp2/3 (Lee *et al*., 2000; Sirotkin *et al*., 2005). However, Myo1-dependent activation of Arp2/3 is dispensable for endocytosis (MacQuarrie *et al*., 2019; Sun *et al*., 2019) and mammalian orthologs, such as the human Myo1e, do not contain CA domains. Additional roles at sites of endocytosis are proposed such as generating force for internalization and/or tethering actin-based processes to the membrane (Sirotkin *et al*., 2005; Berro *et al*., 2010; Basu *et al*., 2014; Lewellyn *et al*., 2015; Sun *et al*., 2019). *Saccharomyces cerevisiae* myosin-1 motor mutants have altered endocytic patch lifetimes (Lewellyn *et al*., 2015; Manenschijn *et al*., 2019; Sun *et al*., 2019), suggesting that Myo1 motor activity is important for its function. In addition, both motor function and TH1 domain membrane binding are required for proper endocytosis (Lewellyn *et al*., 2015; Pedersen and Drubin, 2019). Intriguingly, the majority of both yeast and human long-tailed myosin-1s are found in the cytoplasm (Sirotkin *et al*., 2010; Tanimura *et al*., 2016; Manenschijn *et al*., 2019) but how myosin-1 membrane localization is controlled is unknown.

Here, we identified a previously uncharacterized protein, Ank1, in a Myo1-GFP purification. The ankyrin repeats of Ank1 bind a conserved lever arm residue in Myo1 and the Ank1 acidic tail extends to also bind the Myo1 TH1 domain. This cytoplasmic complex precludes Myo1 from membrane binding and cells lacking this mode of regulation have non-specific Myo1 membrane localization and endocytic defects. An Ank1 ortholog is also present in budding yeast that regulates the localization of both long-tailed myosin-1s, Myo3 and Myo5. Lastly, the human Ank1 ortholog, OSTF1, controls Myo1e localization to endocytic sites in cultured cells indicating that this mechanism of long-tailed myosin-1 regulation is conserved.

## Results and Discussion

The uncharacterized *S. pombe* protein encoded by SPAC105.02c was identified by mass spectrometry in affinity purifications of Myo1-GFP along with known Myo1 associated proteins including the calmodulin-like light chains Cam1 and Cam2, WIP homolog Bbc1, F-BAR protein Cdc15, verprolin homolog Vrp1 and WASp homolog Wsp1 (Fig. 1A) (Toya *et al*., 2001; Carnahan and Gould, 2003; Sirotkin *et al*., 2005; Sammons *et al*., 2011; MacQuarrie *et al*., 2019). The SPAC105.02c ORF encodes a predicted 18 kDa protein containing three ankyrin repeats and an acidic tail that we named Ank1 (<u>Ank</u>yrin repeat-protein <u>1</u>). The Myo1-Ank1 association was validated by co-immunoprecipitation (Fig. 1B). Cam1 and Cam2 also co-immunoprecipitated with Ank1, dependent on Myo1 (Fig. S1A-B). Thus, we investigated Ank1’s potential relationship to Myo1 function.

**Figure 1.**
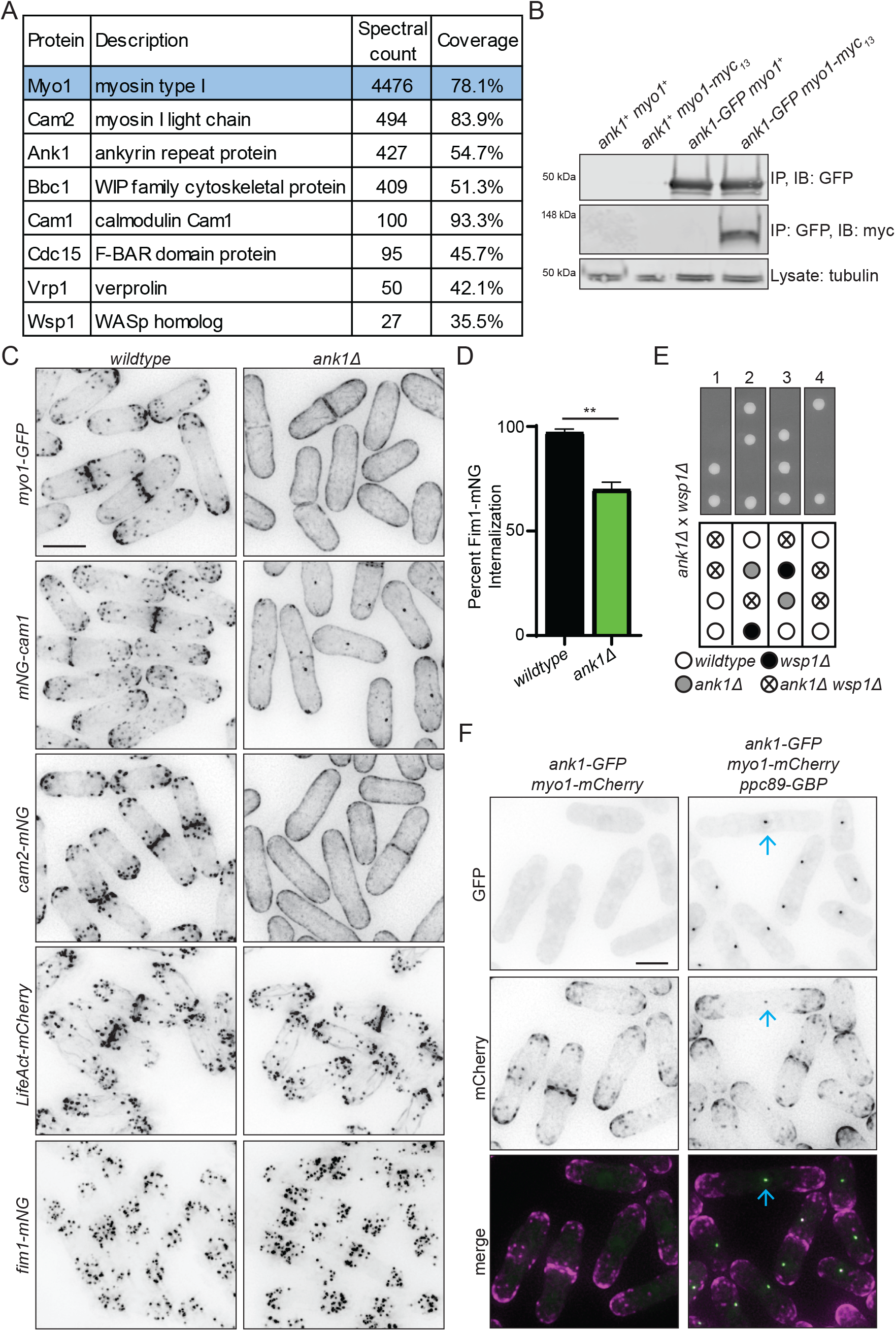
Ank1 associates with and controls the localization of Myo1. A) A summary of data from mass spectrometry analysis of proteins associated with Myo1-GFP. B) Anti-GFP immunoblot (IB; top) and anti-myc immunoblot (middle) of anti-GFP immunoprecipitations (IP) from the indicated strains. α-tubulin was used as a loading control (bottom). Part of this blot is also shown in Figure S1F. C) Representative live-cell images of cells expressing *myo1-GFP, mNG-cam1, cam2-mNG, LifeAct-mCherry* or *fim1-mNG* in wildtype and *ank1Δ* cells grown up at 25°C prior to imaging. D) Quantification of the percent of Fim1-mNG that internalized into the cell in wildtype and *ank1Δ* cells from live-cell time-lapse imaging of the strains imaged with 1s intervals for a total of 1 min. Error bars represent SEM, ** p < 0.01; *t* test. n of patches ≥ 74 from n of cells ≥ 8. E) Representative tetrads and schematic of the indicated genetic cross. F) Live-cell imaging of the indicated strains grown up at 25°C prior to imaging. The blue arrows indicate Ank1-GFP and Myo1-mCherry co-localized at the SPB when Ppc89-GBP is expressed in the cell. Scale bars, 5 µm.

In wildtype cells, Myo1-GFP localized to endocytic actin patches enriched at the cell tips and cell division site, as previously described (Lee *et al*., 2000; Sirotkin *et al*., 2005). In *ank1Δ* cells, Myo1-GFP was instead localized uniformly along the plasma membrane (PM) (Fig. 1C). Because Cam1 and Cam2 directly bind the Myo1 IQ motifs (Toya *et al*., 2001; Sammons *et al*., 2011) and co-immunoprecipitate with Ank1, their localization was also analyzed. Like Myo1-GFP, mNG-Cam1 and Cam2-mNG localized to actin patches in wildtype cells, as previously shown (Sammons *et al*., 2011), but like Myo1, they were also mis-localized along the PM in *ank1Δ* cells (Fig. 1C). mNG-Cam1 also localizes to the cytokinetic ring and spindle pole bodies (SPBs) (Moser *et al*., 1997; Eng *et al*., 1998), and these localizations are unchanged in *ank1Δ* cells (Fig. 1C). F-actin visualized with LifeAct-mCherry and Fim1-mNG localized in patches in wildtype and *ank1Δ* cells indicating that actin patches are present in *ank1Δ* cells (Fig. 1C). Interestingly, although Vrp1 and Wsp1 directly or indirectly associate with Myo1’s C-terminus, respectively (Carnahan and Gould, 2003; Sirotkin *et al*., 2005; MacQuarrie *et al*., 2019), they localized to patches in both wildtype and *ank1Δ* cells (Fig. S1C). Thus, Ank1 specifically controls myosin-1 localization.

To investigate if Ank1 impacts Myo1 function in endocytosis, we analyzed wildtype and *ank1Δ* cells expressing Fim1-mNG. Whereas >95% of Fim1-mNG patches internalized in wildtype cells, as expected (Barger *et al*., 2019), only 65% of patches internalized in *ank1Δ* cells (Fig. 1C-D). Genetic analysis revealed that *ank1Δ* was synthetically lethal with *wsp1Δ* (Fig. 1E) and synthetically sick with *vrp1Δ, as* previously noted (Ryan *et al*., 2012), and was also synthetically sick with *arp3-C1* (Fig. S1D) (McCollum *et al*., 1996). We did not detect a significant genetic interaction between *ank1Δ* and *myo1Δ*, consistent with *ank1* functioning in the same genetic pathway as *myo1* (Fig. S1D). These results implicate Ank1 in promoting Myo1 function in endocytosis.

Because Ank1 co-purified with Myo1, is required for proper Myo1 localization and plays a role in endocytosis, we investigated Ank1’s localization. Unexpectedly, Ank1-GFP localized diffusely in the cytoplasm and did not apparently co-localize with Myo1-mCherry at actin patches (Fig. 1F). However, when Ank1-GFP was targeted to the SPB via Ppc89-GBP (Rothbauer *et al*., 2008), Myo1-mCherry was recruited to the SPB (Fig. 1F and S1E). Taken together, these data support an association between Ank1 with Myo1 and suggest that Ank1 may interact with a pool of Myo1 that is cytoplasmic, an intriguing possibility given 84% of the Myo1 protein is in the cytoplasm (Sirotkin *et al*., 2010).

To further investigate the nature of the Ank1-Myo1 interaction, we performed immunoprecipitations of Myo1 C-terminal truncations (Lee *et al*., 2000) and assayed if Ank1 co-purified. Myo1 lacking the CA [Myo1Δ(1-1165)], CA and SH3 [Myo1Δ(1-1102)], or CA, SH3 and TH2 domains [Myo1Δ(1-964)] still associated with Ank1-GFP (Fig. S1F). In fact, we found that a fragment containing only the motor domain and IQ motifs [Myo1(1-772)] or a shorter fragment containing the motor domain and the first few residues of the lever arm (but not the IQ motifs) [Myo1(1-721)] also still associated with Ank1 (Fig. 2A-B). In accord, Myo1Δ(1-772)-mNG localized to endocytic patches in wildtype cells but in *ank1Δ* cells, Myo1Δ(1-772)-mNG localized along the PM (Fig. 2C), consistent with the Myo1 C-terminal domains being dispensable for Ank1 interaction.

**Figure 2.**
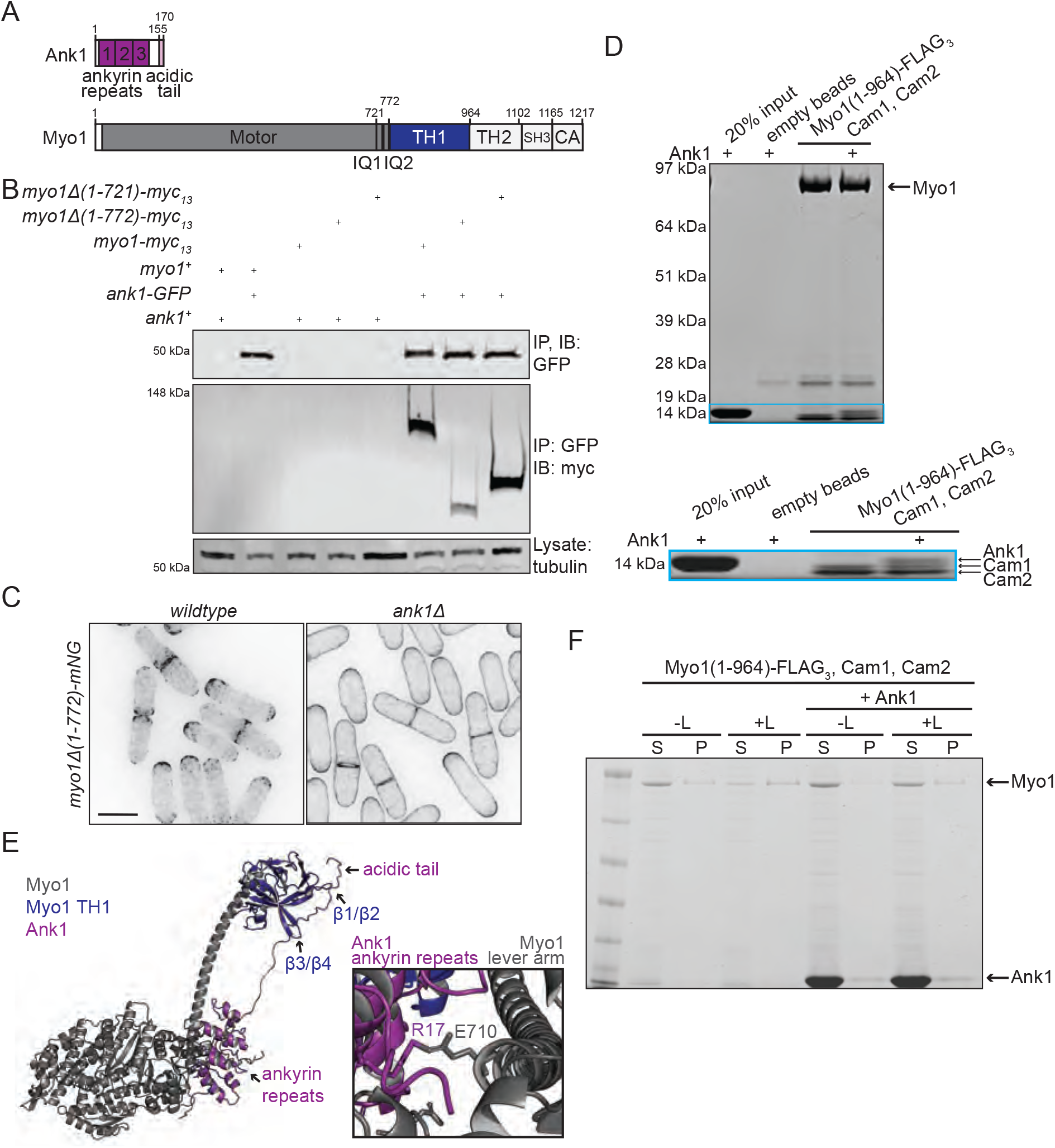
Myo1 and Ank1 are predicted to physically associate. A) A schematic, drawn to scale, of the domain layouts of Ank1 and Myo1 proteins. Myo1 contains the following C-terminal domains: isoleucine-glutamine (IQ), tail homology 1 (TH1), tail homology 2 (TH2), *src* homology 3 (SH3) and central acidic (CA). The numbers on Ank1 indicate the three ankyrin repeats. B) Anti-GFP immunoblot (top) and anti-myc immunoblot (middle) of anti-GFP immunoprecipitations from the indicated strains. α-tubulin was used as a loading control (bottom). C) Live-cell imaging of Myo1Δ(1-964)-mNG in wildtype or *ank1Δ* cells grown up at 25°C prior to imaging. Scale bar, 5 µm. D) In vitro binding assay with bead bound recombinant Myo1(1-964)-FLAG_3_-Cam1-Cam2 with recombinant Ank1. Samples were washed, resolved by SDS-PAGE and stained with Coomassie blue. The second lane is a control with empty beads incubated with Ank1. E) Left, AlphaFold2 (AF) predicted structure of the Myo1-Ank1 complex. The Myo1 motor domain, lever arm and TH1 domain (residues 1-964) were modeled with full-length Ank1. The motor domain and lever arm of Myo1 are gray and the Myo1 TH1 domain is blue. Ank1 is magenta. Right, a zoomed in view of the interaction interface of Ank1 and Myo1 predicted by AF where the ankyrin repeats bind the Myo1 lever arm. F) Co-pelleting between Myo1(1-964)-FLAG_3_-Cam1-Cam2 with and without Ank1 and also Ank1 alone with Folch fraction liposomes (Liposomes = L; supernatant = S, unbound; and pellet = P, bound).

To test for direct binding between Ank1 and Myo1, we co-produced a fragment of Myo1 containing the motor domain, lever arm and TH1 domain tagged with FLAG_3_ [Myo1(1-964)-FLAG_3_] with Cam1 and Cam2 in Sf9 cells and purified the complex. We also purified His_6_-Ank1 from bacteria and removed the tag. The Myo1-Cam1-Cam2 complex bound Ank1, supporting a direct interaction between Myo1 and Ank1 (Fig. 2D).

To gather more information about how Ank1 and Myo1 may associate, we employed AlphaFold2 (AF) to model their interaction (Jumper *et al*., 2021; Mirdita *et al*., 2022; Varadi *et al*., 2022). AF predicted that the ankyrin repeats of Ank1 are positioned at the back of the Myo1 motor domain, opposite from the actin binding cleft, where the lever arm extends (Fig. 2A and E). In addition, the Ank1 acidic tail was predicted to extend away from the ankyrin repeats and bind the Myo1 TH1 domain (Fig. 2E). The myosin-1 TH1 domain is an extended pleckstrin homology domain (Hokanson *et al*., 2006) and typical membrane binding loops (β1/β2 and β3/β4) (Cronin *et al*., 2004; Hokanson *et al*., 2006; Feeser *et al*., 2010; Brzeska *et al*., 2020) is where the Ank1 acidic tail is predicted to associate (Fig. 2E). Given the AF model and the above results, we hypothesized that Ank1 precludes Myo1 membrane binding and sequesters the myosin-1 complex in the cytoplasm (Fig. 2F).

We next tested if Myo1 membrane binding is controlled by Ank1 *in vitro*. As expected, a portion of Myo1(1-964)-FLAG_3_-Cam1-Cam2 co-pelleted with liposomes made from Folch fraction lipids (Fig. 2F). Addition of Ank1 resulted in a reduction in the amount of co-pelleting of the Myo1(1-964)-FLAG_3_-Cam1-Cam2 complex with liposomes (Fig. 2F). We therefore conclude that Ank1 directly binds and inhibits Myo1 membrane binding, supporting our model.

Based on the above results, we predicted that Ank1 overexpression would result in increased cytoplasmic Myo1. In control *myo1-mNG fim1-mCherry* cells, Myo1-mNG and Fim1-mCherry both localized at actin patches (Fig. 3A). In contrast, in *myo1-mNG fim1-mCherry* cells overproducing Ank1, while Fim1-mCherry was at patches, almost all Myo1-mNG was cytoplasmic (Fig. 3A). These data support our model that Ank1 blocks Myo1 membrane association and normally sequesters the bulk of Myo1 in the cytoplasm. Consistent with this conclusion, it was previously shown that overexpressed GFP-Myo1 localized along the entire PM (Lee *et al*., 2000; Attanapola *et al*., 2009), as would be predicted if there was an excess of Myo1 over available Ank1. Furthermore, in cells overproducing *ank1^+^*, ∼10% of Fim1-mCherry patches internalized compared to ∼95% of Fim1-mCherry patches internalized in the control cells (Fig. 3B). These data illustrate the importance of proper regulation of Myo1 membrane localization during endocytosis.

**Figure 3.**
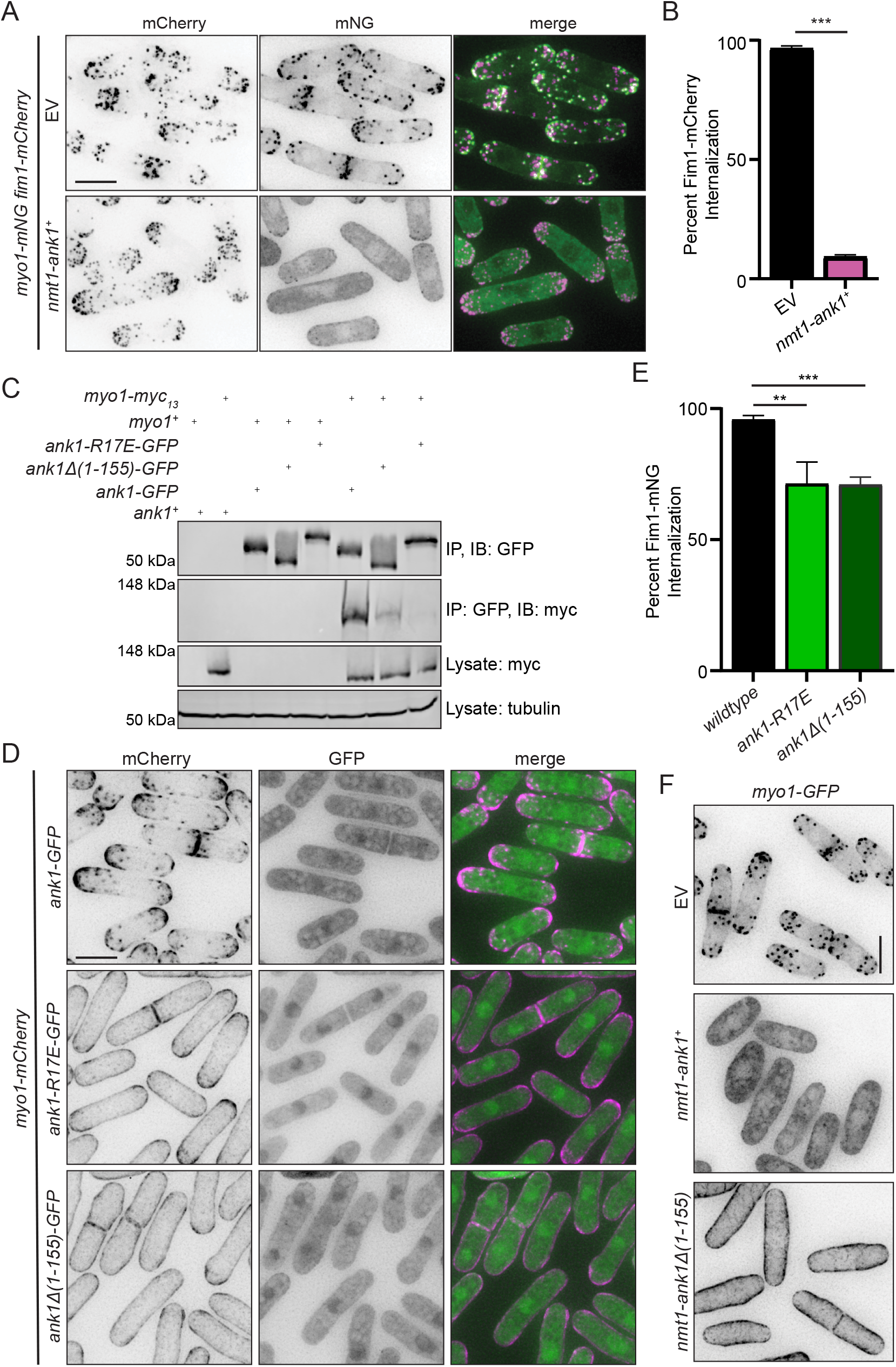
Structure-function analysis of Ank1. A) Live-cell imaging of *myo1-mNG fim1-mCherry* induced to express either an empty vector (EV) or a vector containing *nmt1-ank1^+^* for 24 h at 32°C. B) Quantification of the percent of Fim1-mCherry that internalized into the cell in *myo1-mNG fim1-mCherry* cells expressing EV or *nmt1-ank1^+^* from live-cell time-lapse imaging of the strains from A imaged with 1s intervals for a total of 1 min. Error bars represent SEM, *** p < 0.001; *t* test. n of patches ≥ 85 from n of cells ≥ 9. C) Anti-GFP immunoblot (top) and anti-myc immunoblot (second from top) of anti-GFP immunoprecipitations from the indicated strains. anti-myc immunoblot from lysates is shown in the blot second from the bottom. α-tubulin was used as a loading control (bottom). D) Live-cell imaging of cells expressing Myo1-mCherry with Ank1-GFP, Ank1Δ(1-155)-GFP or Ank1-R17E-GFP. Cells were grown up at 25°C prior to imaging. E) Quantification of the percent of Fim1-mNG that internalized into the cell in wildtype, *ank1-R17E* and *ank1Δ(1-155)* cells from live-cell time-lapse imaging of the strains imaged with 1s intervals for a total of 40s. Error bars represent SEM, ** p < 0.01; *** p < 0.001; *t* test. n of patches ≥ 109 from n of cells ≥ 14. F) Live-cell images of *myo1-GFP* induced to express either an EV, *ank1^+^* or *ank1Δ(1-155)* for 24 h at 32°C. Scale bars, 5 µm.

We next used the AF generated model to inform Ank1 structure-function analysis. AF predicted that Ank1 R17 within the ankyrin repeats binds to Myo1 E710 within the lever arm (Fig. 2D). We therefore constructed an *ank1* allele in which R17 was substituted with glutamate (*ank1-R17E*) to test whether this would disrupt the Myo1-Ank1 interaction. Indeed, Ank1-R17E-GFP did not co-immunoprecipitate with Myo1-myc_13_ (Fig. 3C). Further, Myo1-mCherry localized along the PM in *ank1-R17E-GFP* cells indicating it is a non-functional *ank1* allele (Fig. 3D). In addition, *ank1-R17E* displayed endocytic defects compared to wildtype with only 70% of the Fim1-mNG patches internalizing (Fig. 3E). We conclude that Ank1 R17 is a key residue for binding Myo1.

We next explored the function of the Ank1 acidic tail that AF predicted bound the Myo1 TH1 domain (Fig. 2D). We removed it by adding sequences encoding GFP at the *ank1* endogenous locus to construct the *ank1Δ(1-155)-GFP* strain (Fig. 2A). Immunoprecipitation experiments revealed that Ank1Δ(1-155)-GFP still bound Myo1-myc_13_ but reproducibly less well (Fig. 3C). That the Ank1 acidic tail is not required for Myo1 interaction is consistent with our previous experiment demonstrating that the Myo1 TH1 domain is dispensable for Ank1 co-immunoprecipitation (Fig. 2B). In *ank1Δ(1-155)-GFP* cells, Myo1-mCherry was redistributed all along the PM in a manner indistinguishable from its localization in *ank1Δ* (Fig. 3B). *ank1Δ(1-155)-GFP* cells also displayed endocytic defects with only 70% of Fim1-mNG puncta internalizing (Fig. 3E).

We also assayed the overproduction of Ank1Δ(1-155) in Myo1-GFP cells. Overproduced Ank1Δ(1-155) resulted in Myo1-GFP localizing along the entire PM (Fig. 3F). We interpret these findings to indicate that overproduced Ank1Δ(1-155) can out-compete endogenous Ank1 for binding to Myo1-GFP, but Ank1Δ(1-155) cannot inhibit Myo1’s membrane association, thus resulting in a dominant negative effect. Overall, we conclude that the Ank1 acidic tail negatively regulates Myo1 membrane association.

Because Ank1 appears to be an important Myo1 regulator, we postulated that this mode of control likely exists in other organisms. Indeed, a predicted Ank1 ortholog is encoded by the ORF YGL242C in *Saccharomyces cerevisiae* and we named it Ank1 (Fig. S2A). Large-scale screens found that Ank1 co-purified with both budding yeast myosin-1s, Myo3 and Myo5, and calmodulin Cmd1 (Krogan *et al*., 2006), and that *ank1Δ* has negative genetic interactions with *vrp1Δ, las17Δ* (WASp ortholog) and *arp2-14* (Costanzo *et al*., 2016). We determined that Ank1-mNG is cytoplasmic and does not apparently co-localize with Myo3-mCherry at actin patches (Fig. S2B). In addition, while both Myo3-mNG and Myo5-mNG localize to endocytic patches in wildtype cells (Sun *et al*., 2006), a portion of the myosin-1s were redistributed along the PM in *ank1Δ* cells (Fig. S2C). AF also predicted an interaction between Ank1 and Myo3 or Myo5 in a similar arrangement to Ank1-Myo1 (Fig. S2D). These data indicate that fission and budding yeast Ank1 proteins are likely orthologs.

The budding and fission yeast myosin-1s belong to the long-tailed subclass, a group that is conserved in humans (Myo1e and Myo1f). Thus, we explored if Ank1 orthologs exist beyond yeast. One of the top co-purifying proteins with both human Myo1e and Myo1f was OSTF1 (SH3P2 in mice), a protein with three ankyrin repeats, an acidic tail, an N-terminal proline rich region (PRR) and an SH3 domain (Tanimura *et al*., 2016; Huttlin *et al*., 2017, 2021; Cho *et al*., 2022) (Fig. 4A). Interestingly, it was previously found that OSTF1 and Myo1e interact via two interfaces: the OSTF1 PRR binds the Myo1e SH3 domain and the OSTF1 acidic tail binds the Myo1e TH2 domain (Tanimura *et al*., 2016). Intriguingly, in humans and mice, OSTF1 is cytoplasmic and it interacts with Myo1e in the cytoplasm (Tanimura *et al*., 2016; Nakamura *et al*., 2020). In addition, Myo1e at lamellipodia is also controlled by OSTF1; OSTF1 depletion leads to increased membrane-bound Myo1e and overexpression of OSTF1 results in cytoplasmic Myo1e (Tanimura *et al*., 2016). Because of the striking similarities between OSTF1 and Ank1 in domain structure and function, we used AF to explore if OSTF1 may associate with the Myo1e lever arm and TH1 domain as in the Ank1-Myo1 interaction. AF predicted a multivalent interaction with three interfaces. The first predicted interaction was the previously mapped Myo1e SH3 domain binding to the PRR of OSTF1 (Fig. 4B) (Tanimura et al., 2016). The second predicted interaction was the OSTF1 ankyrin repeats binding near the Myo1e motor domain and lever arm, an interaction not previously reported (Fig. 4B). Lastly, the OSTF1 acidic tail was predicted to bind the Myo1e TH1 domain, and this interaction was not previously noted (Fig. 4B). AF did not predict the reported association between the OSTF1 acidic tail and the Myo1e TH2 domain (Tanimura et al., 2016). Instead, unlike in the Ank1-Myo1 case, AF predicted that the unstructured TH2 domain wraps around the Myo1e motor domain to accommodate the C-terminal Myo1e SH3 domain-OSTF1 PRR interaction (Fig. 4B).

**Figure 4.**
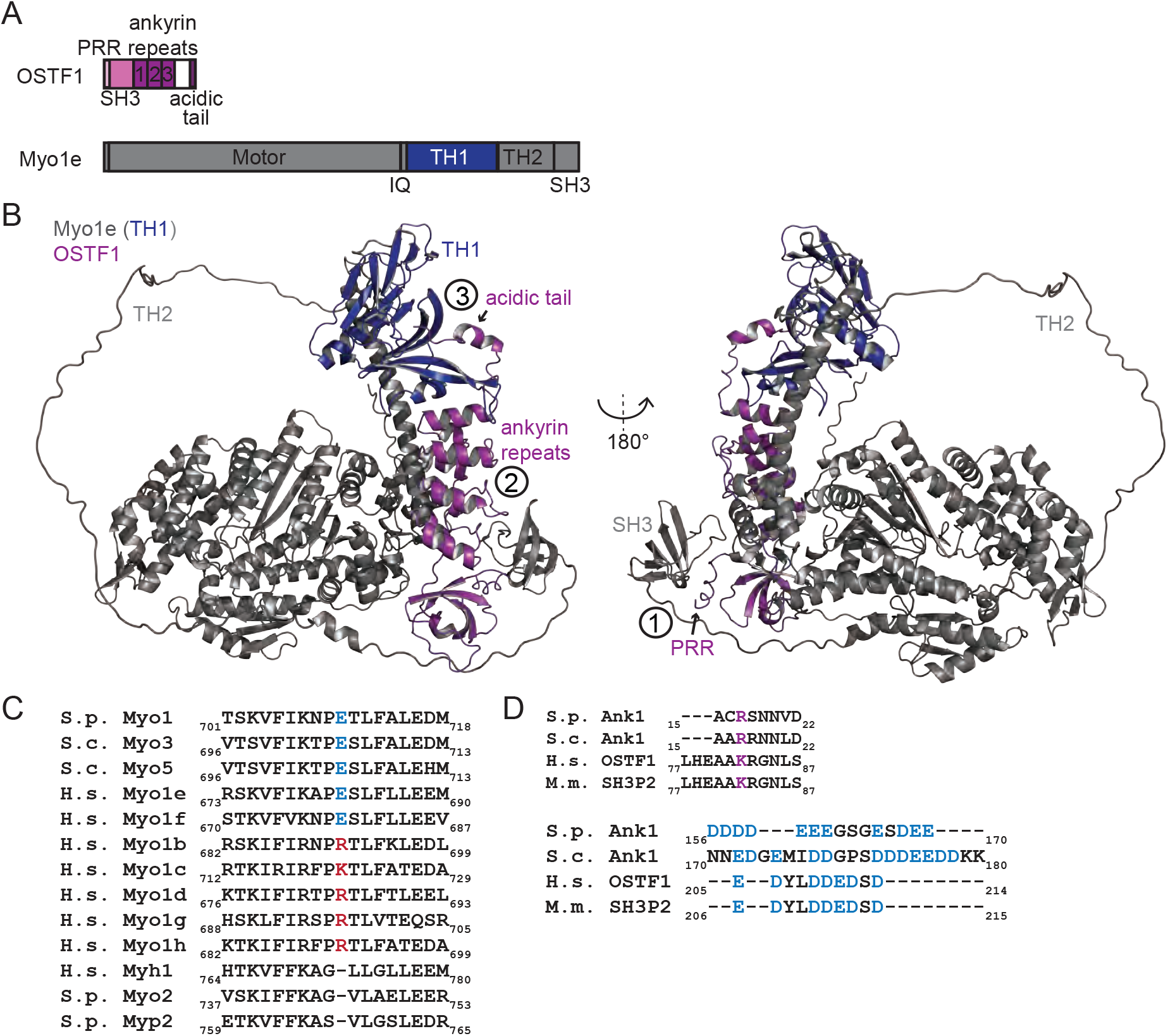
Human OSTF1 is an likely Ank1 ortholog. A) A schematic, drawn to scale, of OSTF1 and Myo1e. Myo1e contains the following C-terminal domains: isoleucine-glutamine (IQ), tail homology 1 (TH1), tail homology 2 (TH2) and *src* homology 3 (SH3). OSTF1 contains an N-terminal proline rich region (PRR), an SH3 domain, three ankyrin repeats and a C-terminal acidic tail. B) AF predicted structure of the OSTF1-Myo1e complex. Full-length proteins were used to model the interaction. Myo1e is in gray except the TH1 domain is blue. The numbers on the model indicate the three predicted interaction interfaces between the proteins. OSTF1 PRR-Myo1e SH3 (1) OSTF1 ankyrin repeats-Myo1e lever arm (2) and OSTF1 acidic tail-Myo1e TH1 (3). OSTF1 is magenta. C-D) Sequence alignments of the indicated amino acids for the indicated proteins. *S. pombe* (S.p.), *S. cerevisiae* (S.c.), *H. sapiens* (H.s.), *M. musculus* (M.m.).

Because of the similarities in predicted structure and that over- or under-expression of OSTF1 modulates Myo1e membrane association (Tanimura *et al*., 2016; Nakamura *et al*., 2020), we hypothesized that these proteins interact in a manner analogous to Ank1-Myo1 to inhibit membrane association of Myo1e and Myo1f. We therefore investigated if the residues involved in Ank1-Myo1 interaction are conserved. The Myo1 glutamate (E710) that is predicted to mediate Ank1 binding is present in all long-tailed isoforms of budding yeast, fission yeast and humans (Fig. 4C). Interestingly, this residue is positively charged or absent in all human short-tailed myosin-1s and all other fission yeast myosins (Fig. 4C). The Ank1 R17 residue necessary for Myo1 interaction is conserved as an arginine or lysine in all Ank1 orthologs identified (Fig. 4D). Lastly, each Ank1 ortholog contains a C-terminal acidic tail of varying length (Fig. 4D). Thus, the Ank1-Myo1 interaction interfaces appear to be conserved.

Depending on the cell type, Myo1e can localizes to a variety of actin structures including lamellipodia, podosomes and sites of endocytosis (Cheng *et al*., 2012; Tanimura *et al*., 2016; Zhang *et al*., 2019). However, OSTF1-dependent control of Myo1e localization has only been demonstrated at lamellipodia (Tanimura *et al*., 2016). To investigate if OSTF1 controls Myo1e endocytic localization we used CRISPR/Cas9-mediated gene editing to insert sequences encoding mNG at the C-terminus of endogenous MYO1E in HeLa cells (Fig. 5A). Correct insertion of the tag was verified by PCR and immunoblotting (Fig. 5A-C). Myo1e-mNG localized to PM puncta in fixed and live cells visualized by total internal reflection fluorescence microscopy (TIRFM) (Fig. 5D-E). The Myo1e-mNG PM puncta represented endocytic sites as they colocalized with fluorescently labeled transferrin (Fig. 5D) and Myo1e membrane puncta were shown previously to be endocytic sites (Cheng *et al*., 2012). Myo1e-mNG cells were transfected with plasmids expressing mCherry or OSTF1-mCherry. In cells overexpressing mCherry, Myo1e-mNG localized to PM puncta normally (Fig. 5E). In contrast, in cells overexpressing OSTF1-mCherry, Myo1e-mNG PM puncta were not detected and the signal was instead diffuse (Fig. 5E). Thus, OSTF1 negatively regulates Myo1e localization to sites of endocytosis and likely represents an Ank1 ortholog.

**Figure 5.**
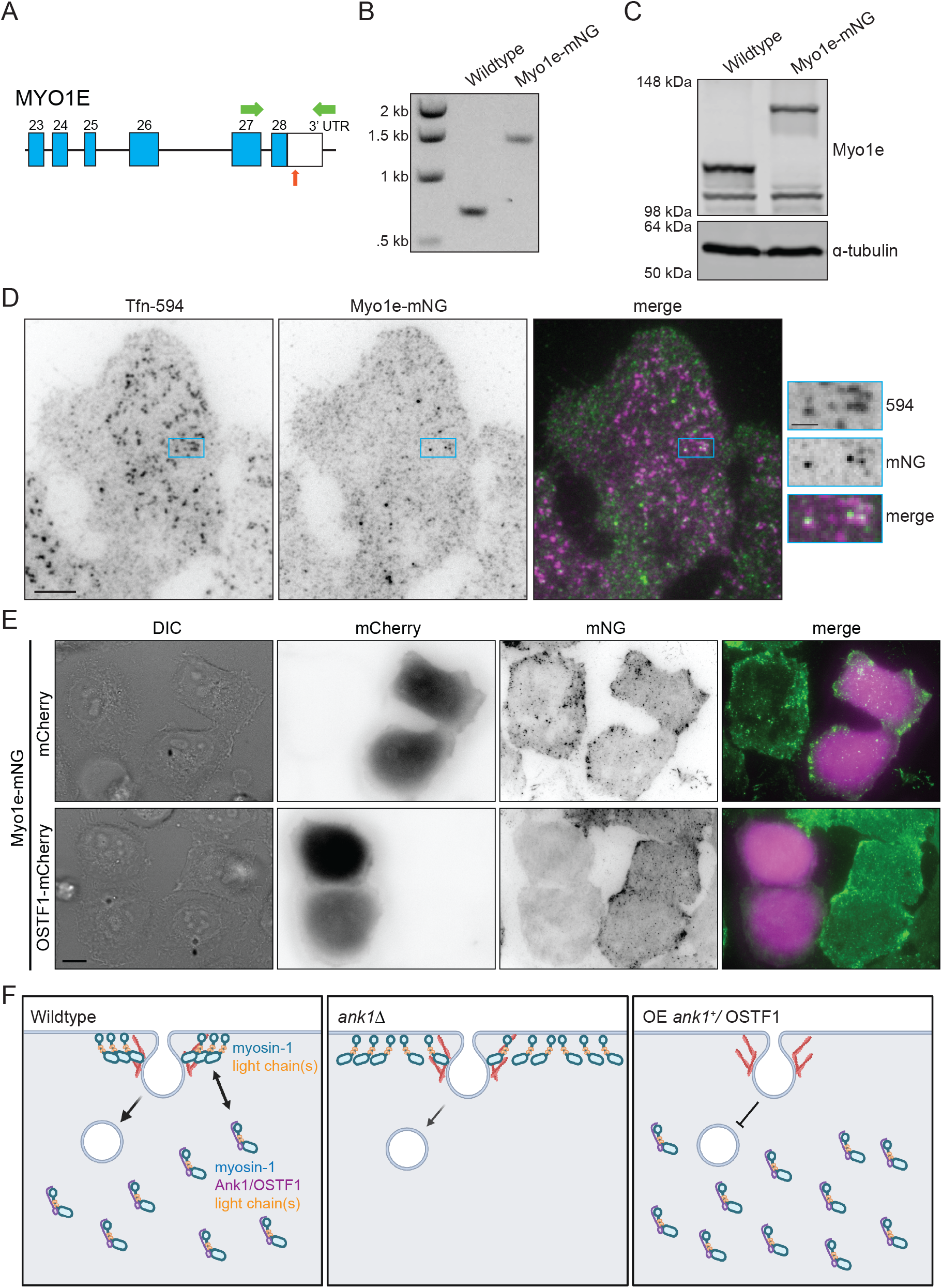
Human OSTF1 controls Myo1e localization at sites of endocytosis. A) Cartoon representation of the 3’ end of the MYO1E gene. The orange arrow points to the position where the gRNA was designed to target Cas9 for gene editing. Numbers correspond to exons. Green arrows indicate positions of primers used to verify insertion of the tag by PCR analysis. B) PCR verification of the cell line with endogenously tagged MYO1E. C) Immunoblot analyses of whole cell lysates from wildtype and Myo1e-mNG cells with anti-α-tubulin (loading control) and anti-Myo1e. D) Fixed-cell TIRFM imaging of Myo1e-mNG cells stained with Alexa Flour 594-transferrin (Tfn-T94). Scale bar on left images, 5 μm and scale bar on zoomed in section on right, 1 μm. E) Live-cell TIRFM of Myo1e-mNG cells expressing mCherry or OSTF1-mCherry. Scale bar, 10 μm. F) In wildtype cells (left) myosin-1 exists in a cytoplasmic pool bound by Ank1/OSTF1, and a membrane bound pool at sites of endocytosis that lack Ank1/OSTF1. When yeast cells lack *ank1^+^* (middle), myosin-1 constitutively binds along the entire PM and these cells have defects in endocytic internalization. Overproduction of Ank1 or OSTF1 (right) causes all the myosin-1 to be cytoplasmic and not localize to sites of endocytosis. In yeast cells this leads to severe endocytic internalization defects. Created with BioRender.com.

Over-production of Ank1 or OSTF1 resulted in a redistribution of the corresponding myosin-1 from endocytic sites to the cytoplasm (Fig. 5F). This suggests that, in addition to inhibiting membrane binding, myosin-1 F-actin binding and/or motor activity may also be inhibited by Ank1/OSTF1. Given where the ankyrin repeats are predicted to bind myosin-1, it is possible that the myosin-1 ATPase cycle is negatively impacted and/or the lever arm is physically constrained. Future studies are required to understand how, in addition to membrane binding, myosin-1 activities are controlled by Ank1/OSTF1.

Conversely, *ank1* deletion results in a milder endocytic defect compared to *myo1Δ* (Basu *et al*., 2014; Barger *et al*., 2019). In *ank1Δ* cells, myosin-1 is uniformly distributed along the PM but not specifically concentrated at sites of endocytosis (Fig. 5F). Therefore, when an actin patch forms, a fraction of myosin-1 may be available to promote endocytosis, potentially explaining the relatively mild *ank1Δ* endocytic defect. The precise role of myosin-1s in endocytosis is still not fully understood. One model proposes that myosin-1 motors translocate F-actin away from the PM to facilitate monomer addition at barbed ends and growth of the F-actin network (Manenschijn *et al*., 2019). By preventing spurious PM localization, Ank1/OSTF1 may promote the local accumulation of myosin-1 at endocytic sites.

One remaining question is how the Ank1/OSTF1-myosin-1 associations are controlled to ensure the proper balance of myosin-1 at membranes and in the cytoplasm. Several regulatory mechanisms have been proposed for OSTF1. First, OSTF1 is phosphorylated by ERK on S202 to control its association with Myo1e (Tanimura *et al*., 2016). Fission yeast Ank1 phosphorylation sites are reported (Swaffer *et al*., 2018), but none are analogous to S202 or in proximity to the predicted interaction interfaces. OSTF1 also has additional binding partners, one of which, RP2, is proposed to regulate the subcellular localization of OSTF1 and its ability to bind and regulate Myo1e (Lyraki et al., 2018). Further studies will be needed to elucidate the full set of regulatory mechanisms affecting Ank1/OSTF1-myosin-1 complex formation.

Although OSTF1 and Ank1 have many similarities in function and protein structure, OSTF1 contains additional N-terminal domains that are not found in the yeast orthologs. One of these, the PRR, binds the Myo1e SH3 domain which is predicted to result in the unstructured TH2 domain of Myo1e looping around the motor domain. The TH2 domain also contributes to Myo1e membrane binding (Zhang *et al*., 2019). Thus, interaction between the OSTF1 PRR and the Myo1e SH3 domain may preclude TH2 from associating with the PM. How exactly each interaction interface influences myosin-1 function will be interesting to explore further.

## Materials and Methods

### Yeast methods

All *S. pombe* strains used in this study (Table S1) were cultured using standard methods in YE media or Edinburgh minimal media plus selective supplements (Moreno *et al*., 1991; Forsburg and Rhind, 2006). All budding yeast strains used in this study (Table S1) were derived from wildtype diploid S288C and cultured in YPD using standard methods. Transformation of yeast with plasmid or linear DNA was accomplished using electroporation (Forsburg and Rhind, 2006) and lithium acetate methods (Keeney and Boeke, 1994; Forsburg and Rhind, 2006), respectively. Strain construction was accomplished through tetrad analysis using standard methods.

Tagged strains were generated by endogenously tagging the 3′ end of ORFs with sequences encoding the appropriate epitope tag or fluorescent protein and/or resistance or auxotrophic selection cassette (*kanMX6, natMX6, or hphMX6*) using pFA6 cassettes as previously described (Wach et al., 1994; Bähler et al., 1998). G418 (Geneticin, 100 mg/mL, Thermo Fisher Scientific; cat# 11811031), hygromycin B (50 mg/mL, Life Technologies Cat# 10687010), or nourseothiricin (clonNAT, 100 mg/mL, GoldBio; cat# N-500-100) were used for selection of *kanMX6, natMX6, or hphMX6* cells, respectively. All fusion proteins were expressed from their native promoters at their chromosomal loci.

The *ank1Δ::ura4^+^* allele was constructed by performing a marker swap from the *ank1::kanMX6* strain (Bioneer). The *ank1* gene was amplified from genomic DNA with 300bp flanking regions by PCR and cloned into BamHI/PstI sites in pIRT2 by Gibson assembly. The *ank1-R17E* allele was created by site-directed mutagenesis and sequenced to confirm. To integrate the *ank1-R17E* allele into the genome, a haploid *ank1Δ::ura4^+^* strain was transformed with pIRT2*-ank1-R17E* and selected on minimal media lacking leucine. Colonies were then grown up in YE and plated on YE plates containing 1.5 mg/mL of 5-fluoroorotic acid (United States Biological; cat# F5050). Integrants were validated by colony PCR and all constructs and integrants were sequenced to ensure their accuracy.

Overexpression of Ank1 was accomplished by using the repressible nmt1 promoter in pREP1. 1 μg of pREP1, pREP1-*ank1^+^* and pREP1-*ank1Δ(1-155)* were transformed into *myo1-mNG fim1-mCherry* or *myo1-GFP* strains by electroporation and plated in medium containing 5 μg/mL thiamine. Cells were then grown in liquid media in the presence of thiamine before being washed into medium lacking thiamine for 24 h to induce overexpression.

### DNA constructs

All cloning was performed by Gibson Assembly. The pSK-*mNG-cam1:kanMX6* construct to make the mNG-Cam1 strain was made by cloning the *cam1* 5’ flank, mNG and *cam1* coding sequence into the NotI/BamHI restriction sites in pSK. Then the sequences encoding kan^R^ and *cam1* 3’ flank were cloned into BamHI/KpnI. *ank1^+^* was amplified from cDNA and cloned into pREP1 and pET15b at the NdeI/BamHI.

The *cam1^+^* coding sequence was cloned into the KpnI/XhoI sites of pFastBac-Dual (Thermo Fisher Scientific; 10712024). The *cam2^+^* coding sequence was then cloned into the BamHI/SacI sites. In addition, the sequence encoding Myo1(1-964)-FLAG_3_ was synthesized in a gene block (Integrated DNA Technologies) and codon optimized for expression in Sf9 cells and cloned into the BamHI/SacI sites in pFastBac-Dual. The *ank1^+^* sequence was cloned in the KpnI/XhoI sites.

pcDNA-OSTF1-mCherry were made by amplifying OSTF1 cDNA from an ORF clone (Sino Biological; cat# HG20537-U) and cloned into the BamHI/EcoRI sites in pcDNA. mCherry was cloned into the EcoRI site of pcDNA.

### Cell culture and gene editing

HeLa cells (TKG cat# TKG 0331, RRID:CVCL_0030) were cultured in Dulbeco’s modified eagle medium (Thermo Fisher Scientific; cat# 11965092) supplemented with 10% fetal bovine serum (Fisher Scientific; cat# 50-152-7066) and 1% penicillin/streptomycin (Thermo Fisher Scientific; cat# 15140122). Cells were plated on glass bottom dishes (MatTek; cat# P35G-1.5-20-C) before transfection with pcDNA-mCherry or pcDNA-OSTF1-mCherry constructs using lipofectamine 3000 reagent (Thermo Fisher Scientific; cat# L3000001).

HeLa cells were fixed with 3.5% PFA for 15 min at RT. Cells were washed with TBS-BSA and then incubated with TBS-BSA and 20 μg/mL Alexa 594-conjugated human transferrin (Thermo Fisher Scientific; cat# T13343) for 1h at RT. Coverslips were washed with TBS-BSA and then mounted on slides using Prolong Diamond antifade mounting media (Thermo Fisher Scientific; cat# P36961).

For CRISPR/Cas9 gene editing, guide RNAs (gRNA) were designed using the “design custom gRNA” tool from Integrated DNA Technologies (IDT) website (https://www.idtdna.com/site/order/designtool/index/CRISPR_CUSTOM). Three gRNAs were chosen based on high on-target and off-target scores. Single guide RNA (sgRNA) containing the gRNA, crRNA, and trans-activating crRNA were synthesized by Synthego Corporation. Tagging *MYO1E* with mNG by CRISPR-mediated gene-editing was performed as described (Paix *et al*., 2017) with modifications. Neon transfection system (Thermo Fisher Scientific; cat# MPK1025) was used for introduction of sgRNA/Cas9 RNP complex into cells. Briefly, 0.5 μL of 15 μM sgRNAs were mixed with 0.5 μL of 15 μM Alt-R HiFi-Cas9 (IDT) in Neon transfection resuspension buffer R (total volume 5 μL) and incubated for 10 min at RT, followed by adding 0.5 μL of 2.8 μM repair template, which is purified PCR product of mNG with 35 bp homologous sequences to either ends of *MYO1E* stop codon. The preformed RNP complex were delivered into 5×10^4^ HeLa cells by electroporation with Neon transfection system using 10 μL Neon tips per manufacturer’s instructions under the condition of voltage 1005 V, width 35 ms and pulse number 2, and the electroporated cells were cultured in 500 μL antibiotics free medium in the presence of 1 μM HDR enhance (IDT) in 24-well plates. Medium was replaced with regular growth medium after 24 h. Three days after electroporation, green positive cells were sorted into single cells in 96-well plates through the Vanderbilt Flow Cytometry core facility. Clones were validated by whole cell PCR amplification of a fragment between 200 bp upstream and 500 bp downstream of *MYO1E* stop codon (with a product of 1.4 kb suggesting successfully tagged clone and that of 700 bp for untagged clone). Edited clones were further verified by Sanger DNA sequencing. Correct edits were also verified by western blotting of whole cell lysate.

### Microscopy

Yeast cells were grown at 25°C prior to live-cell imaging unless otherwise stated. All fission yeast imaging was performed in YE media, except for figures 3A and 3F which were imaged in minimal media supplemented with adenine, uracil and leucine. Budding yeast strains from Figure S2 were grown up in YPD and washed with water prior to imaging. Images of yeast and HeLa cells were acquired with a Personal DeltaVision microscope system (Leica Microsystems) that includes an Olympus IX71 microscope, 60X 1.42 NA PlanApo oil immersion objective, a 60X 1.49 TIRF objective, a pco.edge sCMOS camera, and softWoRx imaging software. TIRF mNG images were acquired with a Sapphire LP 50 25 mW 488 laser (Coherent). Images in figures are deconvolved maximum intensity projections of z sections spaced at 0.5 µm unless otherwise indicated. Quantification of images was performed using Fiji (a version of ImageJ software available at https://fiji.sc) (Schindelin et al., 2012). Percent actin patch internalization was measured by acquiring 40-60s movies with a 1s interval focused at the medial z plan of the cell as determined by the DIC channel. Patches were followed over a 20-second time period in at least 8 cells. Internalization was considered successful if the patch moved at least 2 pixels from the cell membrane. For live-cell imaging of HeLa cells the environmental chamber was set to 37 °C with 5% CO_2_.

### Biochemistry methods

Yeast cell pellets (30 OD) were snap frozen and lysed by bead disruption using a FastPrep-24 cell homogenizer (MP Biomedicals; cat# 116004500) for 20s in NP-40 lysis buffer (6 mM Na_2_HPO_4_, 4 mM NaH_2_PO_4_, 1% NP-40, 150 mM NaCl, 50 mM NaF, 4 μg/ml leupeptin, 0.1 mM Na_3_VO_4_) (Gould et al., 1991) with the addition of 1 mM PMSF, 2 mM benzamidine and protease inhibitor cocktail (Sigma-Aldrich; cat# 11697498001). Buffer modifications were made for the following figures: Figure S1A used NP-40 lysis buffer with 2 mM EGTA and Figure 3C used NP-40 lysis buffer with 300 mM NaCl. For immunoprecipitation, 5 µL of GFP-TRAP magnetic beads (Proteintech; cat# gtma) or mNG-TRAP magnetic beads (Proteintech; cat# ntma) were added to cleared cell lysates, then beads were washed 3 times with NP-40 buffer and proteins were eluted with SDS sample buffer. Proteins were separated on 12%, 10%, 6% Tris-glycine or NuPAGE 4-12% Bis-Tris gels (Invitrogen; cat# NP0321BOX) and transferred to Immobilon-P PVDF membranes (Millipore Sigma; cat# IPVH00005) and immunoblotted with 12CA5 anti-myc (Vanderbilt Antibody and Protein Resource Core) (1:1000), anti-GFP (Sigma-Aldrich; cat# 11814460001, RRID:AB_390913) (1:1000), anti-RFP (Proteintech; cat# 67378-1-Ig, RRID:AB_2882625) (1:1000) or (ChromoTek Cat# 6g6-100, RRID:AB_2631395) (1:2000), anti-mNG (Vanderbilt Antibody and Protein Resource Core, VU527) (1:1000) or anti-ɑ-tubulin (Sigma-Aldrich Cat# T5168, RRID:AB_477579) (1:5000) followed by secondary antibodies conjugated to IRDye 680 or IRDye 800 (LI-COR Biosciences) (1:10000) and imaged with an Odyssey CLx instrument (LI-COR Biosciences).

HeLa cell pellets were lysed in 1% NP-40 buffer containing 6 mM Na_2_HPO_4_, 4 mM NaH_2_PO_4_, 1% NP-40, 150 mM NaCl, 2 mM EDTA, 50 mM NaF, 0.1 mM Na_3_VO_4_ and supplemented with 1 mM PMSF, 2 mM benzamidine and protease inhibitor cocktail. For immunoblotting, whole cell lysate was resolved by SDS-PAGE on a 6 or 12% gel and transferred by electroblotting to PVDF membrane. Proteins were detected with anti-ɑ-tubulin (Thermo Fisher Scientific; cat# MA1-80017, RRID:AB_2210201) or anti-Myo1e (Sigma-Aldrich; cat# HPA023886, RRID:AB_1854253). Immunoblots were Imaged with an Odyssey instrument.

### Protein expression and purification

Rosetta2(DE3)pLysS bacteria cells were grown in Terrific Broth media (23.6 g/L Yeast Extract, 11.8 g/L tryptone, 9.4 g/L K_2_HPO_4_, 2.2 g/L KH_2_PO_4_, 4 mL/L glycerol) with appropriate antibiotics to log-phase (OD_595_ 1–1.5) at 36°C. Cells were incubated on ice for 15 min and protein expression was initiated with the addition of 0.4 mM isopropyl β-D-1-thiogalactopyranoside (IPTG) (Fisher Scientific; BP1755). Cells were then incubated for 16–18 hr at 18°C for optimal protein production.

To purify His_6_-Ank1 from bacteria, frozen cell pellets expressing His_6_-Ank1 were lysed in 20 mM Tris-HCl pH 7.4 and 150 mM NaCl, 1 mM DTT, 1 mM PMSF, 2 mM benzamidine, and protease inhibitor cocktail. Lysates were sonicated three times for 30s, with a 30s pause between sonications (Sonic Dismembrator Model F60, Fisher Scientific; power 15 watts). Lysates were cleared for 15 min at 12k rpm. Cleared lysate was then used in a batch purification protocol by addition of cOmplete His-Tag resin (Roche; 5893682001) for 1 hr at 4°C. Resin was then washed three times with buffer. Protein was eluted with 200 mM imidazole, incubated with thrombin for 2 h at RT and concentrated (Amicon Ultra-0.5 centrifugal filter unit).

Myo1(1-964)-FLAG_3_, Cam1, Cam2 were expressed in baculovirus-infected Sf9 insect cells using the Bac-to-Bac system (Invitrogen). Sf9 cells were coinfected with recombinant baculovirus and after 72 h at 27 °C, the cells were harvested by centrifugation at 5,000 × g for 10 min at 4 °C. The cells were resuspended in ice-cold buffer containing 50 mM Tris-HCl pH 7.4 150 mM NaCl, 2 mM EGTA, 1 mM DTT, 1 mM PMSF, 2 mM benzamidine, and protease inhibitor cocktail and lysed by sonication on ice. Cell debris was pelleted by centrifugation at 12k rpm for 10 min at 4 °C. The clarified supernatant was batch-incubated with Fab-TRAP agarose (Proteintech, ffa) for 1 h at 4°C. The resin was then washed with buffer.

### In vitro binding

In vitro binding assays were performed in buffer containing 20 mM Tris-HCl pH 7.4, 150 mM NaCl, 2 mM EGTA and 1 mM DTT. Recombinant Myo1(1-964)-FLAG_3_-Cam1-Cam2 conjugated to Fab-TRAP beads were incubated with recombinant Ank1 for 1 h at 4°C before washing samples in buffer. Samples were resolved by SDS-PAGE for Coomassie blue staining and imaged on an Odyssey CLx instrument.

### Liposome methods

Liposome were made from Folch fraction lipids (Sigma Aldrich; B1502). CHCl_3_ lipid stocks were mixed and dried under N_2_ gas followed by removal of chloroform under high vacuum. Liposomes were then rehydrated in 50 mM Tris-HCl pH 7.4 and 150 mM NaCl 2 mM EGTA for 30 mins, vigorously vortexed for 1 min and then subjected to ten freeze/thaw cycles and extruded through polycarbonate filters of 400 nm pore size (Millipore Sigma; WHA10417104) using a mini-extruder (Avanti Polar Lipids; 610017).

For co-pelleting assays Myo1(1-964)-FLAG_3_, Cam1, Cam2 with or without 1-2 uM Ank1 was added to 5-10 nM liposomes or buffer for control. The reaction was incubated for 30 min at 4°C and then centrifuged at 150,000 *g* in an Optima TL ultracentrifuge for 15 min at 25°C. Pellet and supernatant fractions were resuspended in equal volumes and analyzed by SDS-PAGE. Protein bands were visualized using Coomassie staining and imaged using an Odyssey CLx instrument.

### Large-scale purifications and LC-MS/MS analysis

A 2 L culture of *myo1-GFP* was grown in 4X YE media (meaning 4x the concentration), generating a pellet of cells (approximately 20 mL packed volume). Cells were lysed under native conditions in a glass bead beater with NP-40 lysis buffer. The lysate was cleared at low speed (3000 rpm) on a tabletop centrifuge. The supernatant was transferred to a clean tube and 30 μL of GFP-TRAP magnetic agarose beads (Proteintech, cat# gtma) pre-washed with NP-40 buffer were added and nutated at 4°C for 1-1.5 h. The beads were then washed with NP-40 buffer two times (5 mL) and then once with 5 mL low-NP-40 buffer (0.02% NP-40) prior to elution from beads with 100 μL of 200 mM glycine twice. The eluate was TCA precipitated (25% TCA on ice) and proteins were identified using liquid chromatography-mass spectrometry (LC-MS) as detailed below. A control purification was also performed from untagged yeast and proteins identified by MS were subtracted from the list of potential interactors.

### MS analysis

Protein pellets were resuspended, digested with trypsin and analyzed by 2D-LC-MS/MS as previously described (Chen et al., 2013) using a Thermo LTQ (Thermo Scientific) with the following modifications: Peptide identifications were filtered and assembled using Scaffold (version 4.4.8; Proteome Software) using the following filters: minimum of 99.0% protein identification probability, minimum of 2 unique peptides, minimum of 95% peptide identification probability. Protein hits were further filtered in Excel to define proteins localized to the division site (TSC ≥ 5) according to Pombase.org (Wood et al., 2012).

### Protein structure prediction and sequence alignment

Protein structure predictions were generated with the ColabFold interface to the AlphaFold2 pipeline on the Colab platform (AlphaFold2.ipynb) (Jumper *et al*., 2021; Mirdita *et al*., 2022; Varadi *et al*., 2022). Protein sequence alignments were performed using Clustal Omega (Madeira *et al*., 2022).

### Statistical analysis

All statistical analysis was performed in Prism 8 (Graphpad software).

## Data availability statement

The data underlying Figs. 1-5 and supplemental Figs. 1-2 are openly available in Mendeley Data at doi: 10.17632/wc4zfr4nrw.1.

## Abbreviations

(MS): mass spectrometry
(mNG): mNeonGreen
(PM): plasma membrane
(TH1): tail homology
(TH2): tail homology 2
(SH3): *src* homology-3
(CA): central acidic
(IQ): Isoleucine-glutamine
(PRR): proline rich region
(AF): AlphaFold2

## Acknowledgements

We thank Jason MacGurn for budding yeast strains and Sierra Cullati and Tony Rossi for critical reading of the manuscript.

This work was supported by NIH grant R35GM131799 to K.L.G.

## Author contributions

A.H. Willet and K.L. Gould conceived the study. A.H. Willet, L. Ren and J.S. Chen performed the experiments. A.H. Willet analyzed all imaging data, L. Ren performed the purification for the mass spectrometry experiment and J.S. Chen processed the sample and analyzed the data for the mass spectrometry experiment. J.S. Chen designed and constructed the Myo1e-mNG CRISPR cell line. A.H. Willet and K.L. Gould wrote the original draft of the manuscript and all authors edited and revised the final manuscript.

## Competing Interests

The authors declare no competing interests.

---

**Supplemental Figure 1 – Ank1 associates with myosin-1.**

**Supplemental Figure 2 – Budding yeast contains an Ank1 ortholog.**

**Supplemental Table 1 – *s. pombe* and *s. cerevisiae* strains used in this study.**

## Notes

### Competing Interest Statement

The authors have declared no competing interest.

