## Supplemental Data 1 for "Membrane binding of endocytic myosin-1s is inhibited by a class of ankyrin repeat proteins"

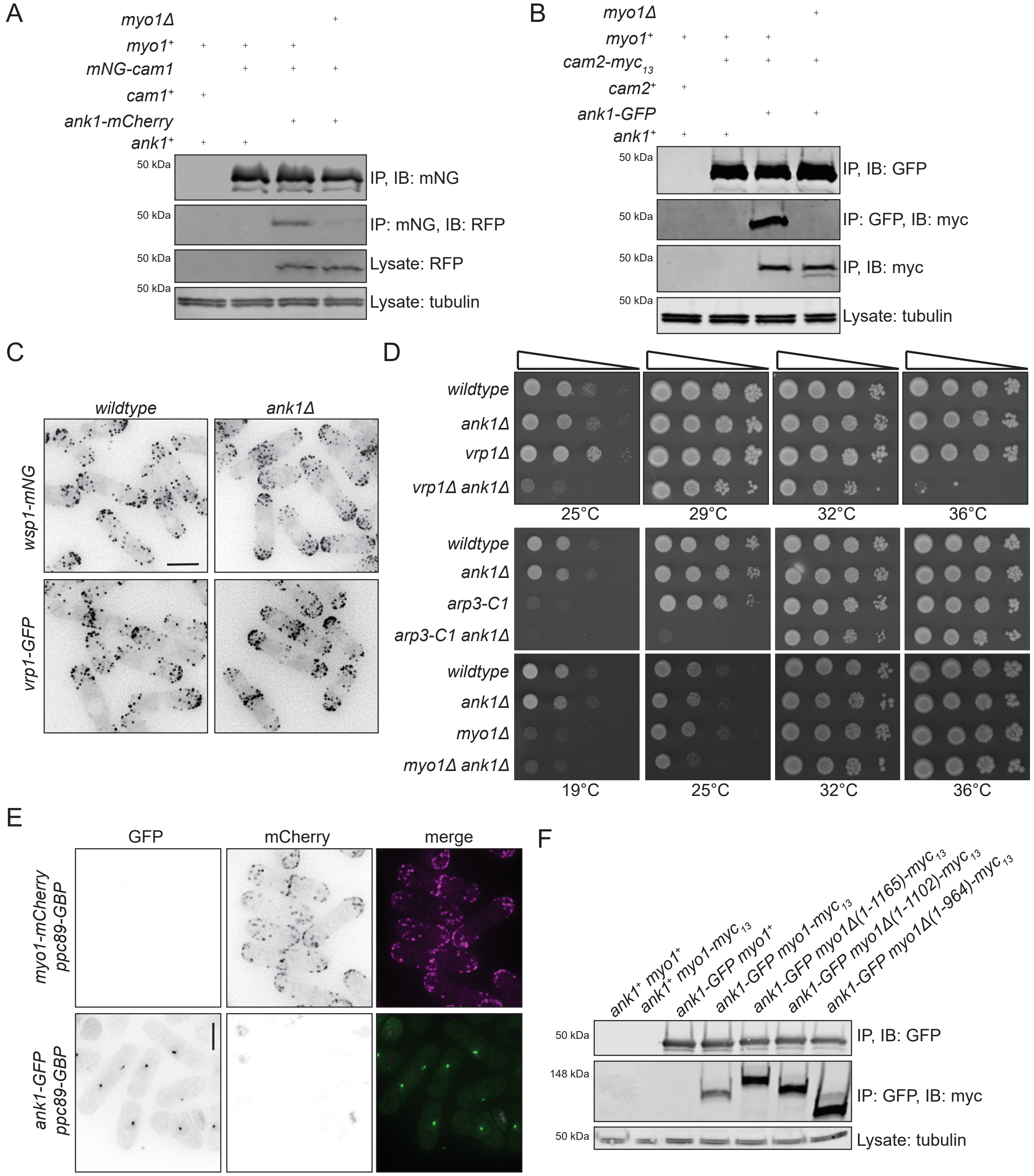

A
myo1Δ
myo1+
mNG-cam1
cam1+
ank1-mCherry
ank1+
50 kDa
IP, IB: mNG
IP: mNG, IB: RFP
Lysate: RFP
Lysate: tubulin
B
myo1Δ
myo1+
cam2-myc13
cam2+
ank1-GFP
ank1+
50 kDa
IP, IB: GFP
IP: GFP, IB: myc
IP, IB: myc
Lysate: tubulin
C
wildtype
ank1Δ
wsp1-mNG
vrp1-GFP
D
wildtype
ank1Δ
vrp1Δ
vrp1Δ ank1Δ
25°C
29°C
32°C
36°C
wildtype
ank1Δ
arp3-C1
arp3-C1 ank1Δ
wildtype
ank1Δ
myo1Δ
myo1Δ ank1Δ
19°C
25°C
32°C
36°C
E
GFP
mCherry
merge
myo1-mCherry ppc89-GBP
ank1-GFP ppc89-GBP
F
ank1+ myo1+
ank1+ myo1-myc13
ank1-GFP myo1+
ank1-GFP myo1-myc13
ank1-GFP myo1Δ(1-1165)-myc13
ank1-GFP myo1Δ(1-1102)-myc13
ank1-GFP myo1Δ(1-964)-myc13
50 kDa
148 kDa
IP, IB: GFP
IP: GFP, IB: myc
Lysate: tubulin


Figure S1

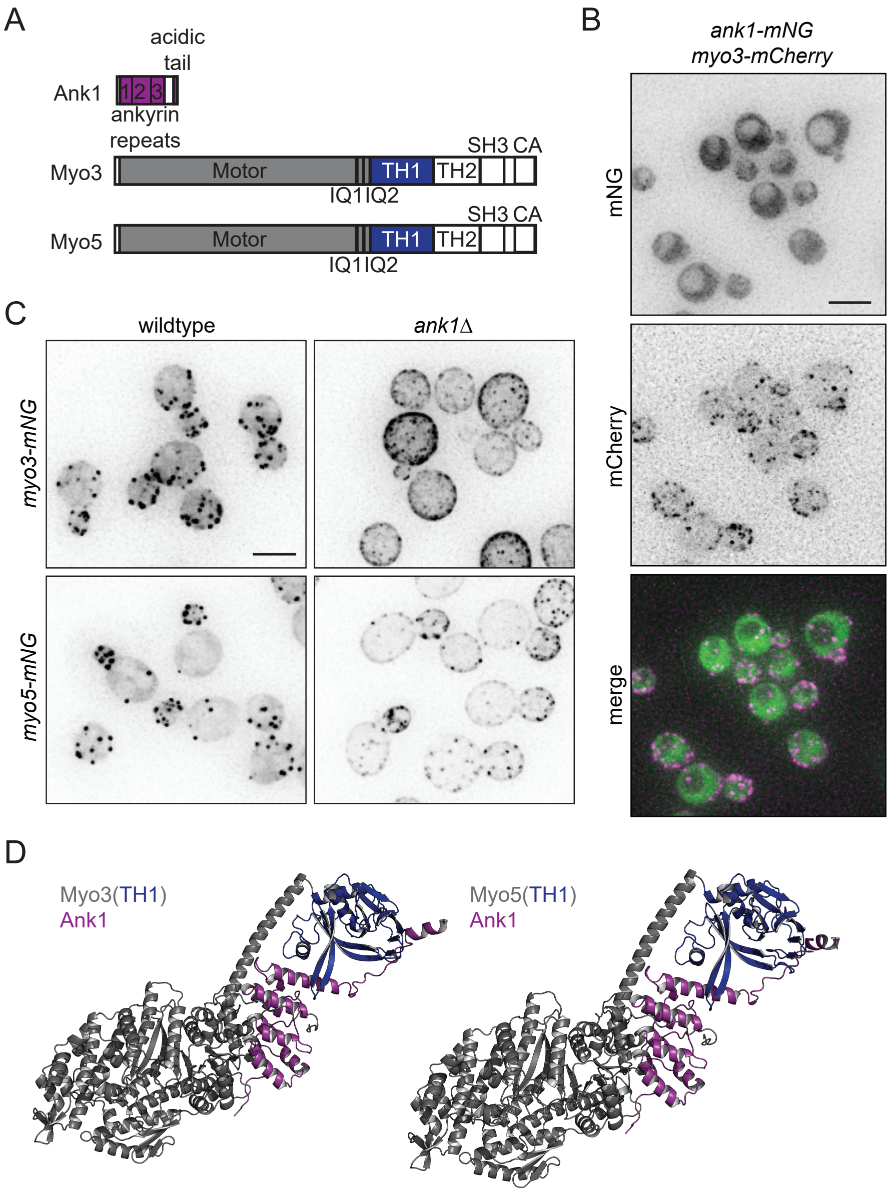
A
acidic tail
Ank1
1 2 3
ankyrin repeats
Myo3
Motor
TH1
TH2
SH3 CA
IQ1 IQ2
Myo5
Motor
TH1
TH2
SH3 CA
IQ1 IQ2
B
ank1-mNG
myo3-mCherry
mNG
mCherry
merge
C
wildtype
ank1Δ
myo3-mNG
myo5-mNG
D
Myo3(TH1)
Ank1
Myo5(TH1)
Ank1

Figure S2

**Supplemental figure 1. Analysis of Ank1 associations and *ank1Δ* cells.** A) Anti-mNG immunoblot (IB; top) and anti-RFP immunoblot (second from top) of anti-mNG immunoprecipitations from the indicated strains. An anti-RFP immunoblot from lysates is second from the bottom and α-tubulin was used as a loading control (bottom). B) Anti-GFP immunoblot (IB; top) and anti-myc immunoblot (second from top) of anti-GFP immunoprecipitations from the indicated strains. Anti-myc immunoblot from lysate samples are shown (second from the bottom) and α-tubulin was used as a loading control (bottom). C) Live-cell images of cells expressing *wsp1-mNG* or *vrp1-GFP* in wildtype and *ank1Δ* cells grown up at 25°C before imaging. D) Serial 10-fold dilutions of the indicated strains spotted on YE agar and incubated for 3-4 d at the indicated temperatures prior to imaging. E) Live-cell imaging of the indicated strains grown up at 25°C prior to imaging. F) Anti-GFP immunoblot (IB; top) and anti-myc immunoblot (middle) of anti-GFP immunoprecipitations from the indicated strains. α-tubulin was used as a loading control (bottom). Part of this blot is also sown in Figure 1B. Scale bars, 5 µm.

**Supplemental figure 2. Budding yeast contains a potential Ank1 ortholog.** A) Schematic, drawn to scale, of the domain layouts of budding yeast Ank1, Myo3 and Myo5 proteins. Myo3 and Myo5 contains the following C-terminal domains: isoleucine-glutamine (IQ), tail homology 1 (TH1), tail homology 2 (TH2), *src* homology 3 (SH3) and central acidic (CA). B) Live-cell imaging of *myo3-mCherry ank1-mNG* cells grown up at 25°C before imaging. C) Live-cell imaging of wildtype and *ank1Δ* cells expressing Myo3-mNG or Myo5-mNG. Cells were grown up at 25°C before imaging. D) Left, AF predicted structure of the Myo3-Ank1 complex. Right, AF predicted structure of Myo5-Ank1 complex. The Myo3/5 motor domain, lever arm and TH1 domain were modeled with full-length Ank1. The motor domains and lever arms of Myo3/5 are gray and the TH1 domains are blue. Ank1 is magenta. Scale bars, 5 µm.

### TABLE S1 *S. pombe* and *S. cerevisiae* strains used in this study

| | | |
|---|---|---|
| **Figure 1** | | |
| KGY3958 | *myo1-GFP:kanMX6 ade6-M21X leu1-32 ura4-D18* $h^-$ | Lab stock |
| KGY246 | *ade6-M210 ura4-D18 leu1-32* $h^-$ | Lab stock |
| KGY19515 | *ank1-GFP:kanMX6 ade6-M210 leu1-32 ura4-D18* $h^-$ | This study |
| KGY3959 | *myo1-*$myc_{13}$*:kanMX6 ade6-M216 leu1-32 ura4-D18* $h^-$ | Lab stock |
| KGY19621 | *ank1-GFP:kanMX6 myo1-*$myc_{13}$*:kanMX6 ade6-M216 leu1-32 ura4-D18* $h^-$ | This study |
| KGY19549 | *myo1-GFP:kanMX6 ank1::kanMX6 ade6-M21X leu1-32 ura4-D18* $h^+$ | This study |
| KGY1944-2 | *mNG-cam1:kanMX6 ade6-M210 leu1-32 ura4-D18* $h^+$ | This study |
| KGY19744 | *mNG-cam1:kanMX6 ank1Δ::kanMX6 ade6-M210 leu1-32 ura4-D18* $h^-$ | This study |
| KGY19766 | *cam2-mNG:hphMX6 ade6-M210 leu1-32 ura4-D18* $h^+$ | This study |
| KGY19767 | *cam2-mNG:hphMX6 ank1Δ::kanMX6 ade6-M210 leu1-32 ura4-D18* $h^+$ | This study |
| KGY14568 | *Pact1-LifeAct-mCherry:*$leu1^+$ *ade6-M210 leu1-32 ura4-D18* $h^+$ | Lab stock |
| KGY19697 | *Pact1-LifeAct-mCherry:*$leu1^+$ *ank1Δ::kanMX6 ade6-M210 leu1-32 ura4-D18* $h^+$ | This study |
| KGY3530-2 | *fim1-mNG:kanMX6 ade6-M210 leu1-32 ura4-D18* $h^+$ | This study |
| KGY19582 | *fim1-mNG:kanMX6 ank1Δ::kanMX6 ade6-M210 leu1-32 ura4-D18* $h^+$ | This study |
| KGY7264 | *wsp1Δ::*$ura4^+$ *ade6-M210 leu1-32 ura4-D18* $h^-$ | Lab stock |
| KGY19489 | *ank1Δ::kanMX6 ade6-M210 leu1-32 ura4-D18* $h^+$ | Bioneer V3 |
| KGY19722 | *ank1-GFP:kanMX6 myo1-mCherry:kanMX6 ade6-M21X leu1-32 ura4-D18* $h^-$ | This study |
| KGY2496-3 | *ank1-GFP:kanMX6 myo1-mCherry:kanMX6 ppc89-GBP:kanMX6 ade6-M21X leu1-32 ura4-D18* $h^-$ | This study |
| **Figure 2** | | |
| KGY148-2 | *myo1Δ(1-721)-*$myc_{13}$*:kanMX6 ade6-M210 leu1-32 ura4-D18* $h^-$ | This study |
| KGY19763 | *myo1Δ(1-772)-*$myc_{13}$*:hphMX6 ade6-M210 leu1-32 ura4-D18* $h^-$ | This study |
| KGY19765 | *ank1-GFP:kanMX6 myo1Δ(1-721)-*$myc_{13}$*:hphMX6 ade6-M21X leu1-32 ura4-D18* $h^-$ | This study |
| KGY4647-2 | *ank1-GFP:kanMX6 myo1Δ(1-772)-*$myc_{13}$*:hphMX6 ade6-M21X leu1-32 ura4-D18* $h^-$ | This study |
| KGY17257 | *myo1Δ(1-964)-mNG:kanMX6 ade6-M210 leu1-32 ura4-D18* $h^-$ | This study |
| KGY19699 | *myo1Δ(1-964)-mNG:hphMX6 ank1Δ::kanMX6 ade6-M210 leu1-32 ura4-D18* $h^+$ | This study |
| **Figure 3** | | |
| KGY4417-2 | *myo1-mNG:hphMX6 fim1-mCherry:natMX6 ade6-M210 leu1-32 ura4-D18* $h^-$ | This study |
| KGY6822-2 | *ank1-R17E-GFP:kanMX6 ade6-M210 leu1-32 ura4-D18* $h^+$ | This study |
| KGY19730 | *ank1Δ(1-155)-GFP:kanMX6 ade6-M210 leu1-32 ura4-D18* $h^-$ | This study |

| | | |
|---|---|---|
| KGY6841-2 | myo1-myc$_{13}$:kanMX6 ank1-R17E-GFP:kanMX6 ade6-M210 leu1-32 ura4-D18 $h^{+}$ | This study |
| KGY19788 | myo1-myc$_{13}$:kanMX6 ank1Δ(1-155)-GFP:kanMX6 ade6-M210 leu1-32 ura4-D18 $h^{+}$ | This study |
| KGY19722 | myo1-mCherry:kanMX6 ank1-GFP:kanMX6 ade6-M210 leu1-32 ura4-D18 $h^{-}$ | This study |
| KGY6840 | myo1-mCherry:kanMX6 ank1-R17E-GFP:kanMX6 ade6-M210 leu1-32 ura4-D18 $h^{-}$ | This study |
| KGY19743 | myo1-mCherry:kanMX6 ank1Δ(1-155)-GFP:kanMX6 ade6-M210 leu1-32 ura4-D18 $h^{+}$ | This study |
| KGY6503-2 | fim1-mNG:kanMX6 ank1-R17E ade6-M210 leu1-32 ura4-D18 $h^{+}$ | This study |
| KGY7198-2 | fim1-mNG:kanMX6 ank1Δ(1-155)-mCherry:natMX6 ade6-M210 leu1-32 ura4-D18 $h^{+}$ | This study |
| **Figure S1** | | |
| KGY7235-2 | mNG-Cam1:kanMX6 ank1-mCherry:natMX6 ade6-M210 leu1-32 ura4-D18 $h^{-}$ | This study |
| KGY7236-2 | mNG-Cam1:kanMX6 ank1-mCherry:natMX6 myo1::kanMX6 ade6-M210 leu1-32 ura4-D18 $h^{+}$ | This study |
| KGY19857 | cam2-myc$_{13}$:hphMX6 ade6-M21X leu1-32 ura4-D18 $h^{+}$ | This study |
| KGY19862 | cam2-myc$_{13}$:hphMX6 ank1-GFP:kanMX6 ade6-M21X leu1-32 ura4-D18 $h^{-}$ | This study |
| KGY19884 | cam2-myc$_{13}$:hphMX6 ank1-GFP:kanMX6 myo1Δ::kanMX6 ade6-M21X leu1-32 ura4-D18 $h^{+}$ | This study |
| KGY18999 | wsp1-mNG:hphMX6 ade6-M210 leu1-32 ura4-D18 $h^{-}$ | This study |
| KGY19700 | wsp1-mNG:hphMX6 ank1Δ::kanMX6 ade6-M210 leu1-32 ura4-D18 $h^{+}$ | This study |
| KGY5664 | vrp1-GFP:kanMX6 ade6-M21X leu1-32 ura4-D18 $h^{+}$ | Lab stock |
| KGY19704 | vrp1-GFP:kanMX6 ank1Δ::kanMX6 ade6-M21X leu1-32 ura4-D18 $h^{+}$ | This study |
| KGY755 | vrp1Δ::ura4$^{+}$ ade6-M210 leu1-32 ura4-D18 $h^{-}$ | Lab stock |
| KGY19646 | ank1Δ::kanMX6 vrp1Δ::ura4$^{+}$ ade6-M210 leu1-32 ura4-D18 $h^{-}$ | This study |
| KGY978 | arp3-C1 ade6-M210 leu1-32 ura4-D18 $h^{-}$ | Lab stock |
| KGY4504-2 | ank1Δ::kanMX6 arp3-C1 ade6-M210 leu1-32 ura4-D18 $h^{+}$ | This study |
| KGY6663 | myo1Δ::kanMX6 ade6-M21X leu1-32 ura4-D18 $h^{-}$ | Lab stock |
| KGY19518 | myo1Δ::kanMX6 ank1Δ::kanMX6 ade6-M21X leu1-32 ura4-D18 $h^{-}$ | This study |
| KGY1897-2 | myo1-mCherry:kanMX6 ppc89-GBP:kanMX6 ade6-M21X leu1-32 ura4-D18 $h^{-}$ | This study |
| KGY233-3 | ank1-GFP:hphMX6 ppc89-GBP:kanMX6 ade6-M21X leu1-32 ura4-D18 $h^{-}$ | This study |
| KGY19677 | ank1-GFP:kanMX6 myo1Δ(1-1165)-myc$_{13}$:hphMX6 ade6-M21X leu1-32 ura4-D18 $h^{-}$ | This study |
| KGY19678 | ank1-GFP:kanMX6 myo1Δ(1-1102)-myc$_{13}$:hphMX6 ade6-M21X leu1-32 ura4-D18 $h^{-}$ | This study |

| | | |
|---|---|---|
| KGY19679 | *ank1-GFP:kanMX6 myo1Δ(1-964)-myc*$_{13}$*:hphMX6 ade6-M21X leu1-32 ura4-D18 h*$^{-}$ | This study |
| **Figure S2** | | |
| KGY5295-2 | *ank1-mNG:hphMX6 myo3-mCherry:natMX6 his3Δ1 leu2Δ0 met15Δ0 ura3Δ0 MATa* | This study |
| KGY5170-2 | *myo3-mNG:hphMX6 his3Δ1 leu2Δ0 met15Δ0 ura3Δ0 MATa* | This study |
| KGY5199-2 | *myo3-mNG:hphMX6 ank1Δ::kanMX6 his3Δ1 leu2Δ0 met15Δ0 ura3Δ0 MATa* | This study |
| KGY5180-2 | *myo5-mNG:hphMX6 his3Δ1 leu2Δ0 met15Δ0 ura3Δ0 MATa* | This study |
| KGY5182-2 | *myo5-mNG:hphMX6 ank1Δ::kanMX6 his3Δ1 leu2Δ0 met15Δ0 ura3Δ0 MATa* | This study |
